# The influence of *Janthinobacterium* sp. strain SLB01 on the cell culture of primmorphs of the sponge *Lubomirskia baicalensis*

**DOI:** 10.1101/2021.12.13.472427

**Authors:** Lubov I. Chernogor, Marina G. Eliseikina, Ivan S. Petrushin, Ekaterina A. Chernogor, Igor V. Khanaev, Sergei I. Belikov

## Abstract

Sponges (phylum Porifera) are ancient, filtering, multicellular metazoans. Freshwater sponges (Demosponges, Lubomirskiidae) dominate the fauna of the littoral zone of Lake Baikal. Over the last years, there have been mass diseases and death of endemic sponges. Previously, the strain *Janthinobacterium* sp. SLB01 was isolated from the diseased sponge *Lubomirskia baicalensis*. The studies of the pathogenicity of the strain *Janthinobacterium* sp. SLB01 for Baikal sponges has not been carried out. Therefore we infected experimentally *in vitro* to determine its pathogenicity by the cell culture of the primmorphs with subsequent isolation, sequencing, and analysis of the genomes. The purpose of the study is to show that the strain *Janthinobacterium* sp. SLB01 isolated from the diseased sponge *L. baicalensis* is a pathogen for the cell culture of primmorphs. The bacteria from the infected samples of the primmorphs were isolated and identified as strain *Janthinobacterium* sp. PLB02. A comparative analysis of the genomes of the strains *Janthinobacterium* sp. SLB01 and *Janthinobacterium* sp. PLB02 showed that they are practically identical. The genomes of both strains contain genes vioABCDE violacein, flok formation, and strong biofilm, and the type VI secretion system (T6SS), as the primary virulence factor. The bacterial strains *Janthinobacterium* sp. based on a comparison of complete genomes showed similarity with strain *Janthinobacterium lividum* MTR. Isolated our strains of *Janthinobacterium* sp. are pathogens for cell cultures of primmorphs and *L. baicalensis* sponges. The results of the study will help to expand the understanding of microbial relationships in the development of disease and the death of Baikal sponges.

## INTRODUCTION

Sponges (phylum Porifera) are ancient, multicellular metazoans (Hedges et al., 2004). They are sessile, filtering invertebrates living in marine and freshwater waters. These multicellular organisms contain prokaryotic and eukaryotic symbiotic microorganisms, including viruses, archaea, cyanobacteria, microalgae, fungi, and protozoa (Osinga et al., 2001; Hentschel et al., 2003; Fieseler et al., 2004; Taylor et al., 2007; Weisz et al., 2008; Webster and Taylor, 2012). Freshwater sponges (Demosponges, Lubomirskiidae) dominate the fauna of the littoral zone of Lake Baikal. Endemic Baikal sponges live in symbiosis with various species of eukaryotes and prokaryotes. Sponges are chlorophyll-containing freshwater organisms due to their association with various chlorophyll-producing algae (Latyshev et al., 1992; Bil et al., 1999; Chernogor et al., 2013). Over the last years, there have been mass diseases and the death of endemic sponges in Lake Baikal (Kravtsova et al., 2014; Khanaev et al., 2018; Belikov et al., 2019). The number of *L. baicalensis* sponges that are most susceptible to the disease has decreased significantly. Annually, up to 10—20% of diseased sponges are observed to die during the winter period (Khanaev et al., 2018). Currently, diseased and dying sponges have been observed in many lake areas (Kravtsova et al., 2014; Khanaev et al., 2018; Belikov et al., 2019). Noticed a large-scale disturbance in the spatial distribution and structure of phytocoenoses of the coastal zone of Lake Baikal (Timoshkin et al., 2016). The etiology and ecology of disease and mass death of sponges remain unknown.

Previously, we showed that in diseased sponges and infected cell culture of the primmorphs occurs mass mortality of green symbionts (Chlorophyta) and a shift in the microbial communities of sponges/primmorphs have been discovered (Belikov et al., 2019; Chernogor et al., 2020). A significant increase in the number of opportunistic microorganisms that belonged mainly to the *Bacteroidetes* and *Proteobacteria*, including *Betaproteobacteria* of the *Oxalobacteraceae* family, was detected using the 16S rRNA gene (Belikov et al., 2019; Chernogor et al., 2020). We isolated the strain *Janthinobacterium* sp. SLB01 bacterium from the diseased sponge *L. baicalensis* collected on the littoral zone of Lake Baikal located in Central Siberia, Russia. The strain *Janthinobacterium* sp. SLB01 of the *Oxalobacteraceae* family has been cultivated, identified as *Janthinobacterium* sp. SLB01, and the draft genome has been published (Petrushin et al., 2019). The strain *Janthinobacterium* sp. SLB01 is a psychrotolerant, violacein–producing, rod-shaped, Gram-negative, aerobic bacterium that contains violacein pigment and can form floc (Petrushin et al., 2020). We identified five genes encoding the VioA, VioB, VioC, VioD and VioE of violacein biosynthesis, which may be a pathogenic factor in this bacterium, similar to the identified genes in the published *J. lividum* MTR strain (Petrushin et al., 2020). It is known that the bacteria produces pigment violacein, which is a compound with antibiotic and antiviral properties (Durán et al., 2007; Rodrigues et al., 2012). In addition, known that the genus *Janthinobacterium* has a wide occurrence ranging from soil, aquatic sites, marine habitats, high altitude environments with a unique ability to survive and colonize new environments (Garrity et al., 2005; Gillis and De Ley, 2006; Gillis and Logan, 2015). We found in this strain key genes of the type VI secretion system (T6SS), which is considered a virulence factor in many *Proteobacteria* (Bingle et al., 2008). Presumably, that genes for the floc formation, including the operon of biosynthesis of exopolysaccharides and an operon containing genes of PEP-CTERM/XrtA system glycosyltransferase, PEP-CTERM system histidine kinase PrsK and PEP-CTERM-box response regulator transcription factor prsR are homologous to the described genes for floc formation in *Zoogloea resiniphila* (Gao et al., 2018).

The studies of the pathogenicity of the strain *Janthinobacterium* sp. SLB01 for Baikal sponges has not been carried out, therefore we experimentally infected cell culture of the primmorphs *in vitro* to determine *Janthinobacterium* sp. strain pathogenicity. The purpose of the study is to show that the strain *Janthinobacterium* sp. SLB01 isolated from the diseased sponge *L. baicalensis* is a pathogen for the cell culture of primmorphs. This research will help clarify one of the reasons for the mass death of Baikal sponges.

## MATERIALS AND METHODS

### Sampling of sponge and cell culture of primmorphs

Specimens of the healthy Baikal sponge *L. baicalensis* Pallas, 1776 (Demospongiae, Haplosclerida, Lubomirskiidae) were collected in individual containers from Lake Baikal in the Olkhon Gate Strait, Russia (53° 02’ 21” N; 106° 57’ 37” E) at a depth of 10 m (water temperature 3–4°C) by scuba divers. The collected samples of green color sponges were transported to the laboratory. The cell cultures of primmorphs were obtained by a method of mechanical dissociation of cells according to the previously described technique (Chernogor et al., 2011). A clean sponge was squashed, and the cell suspension obtained was subsequently filtered through a sterile 200-, 100-, and 29-μm nylon mesh to eliminate pieces of skeleton and spicules of the maternal sponge. The gel-like suspension was diluted 10-fold with Baikal water, placed in a refrigerator, and stored for 3 min at 3–6 °C until a dense precipitate formed. The healthy primmorphs were placed into 200–500-ml cultural bottles (Nalge Nunc International, Rochester, NY, USA). Cell cultures of the primmorphs were cultivated in NBW at 3–4°C and light intensity of 47 lux or 0.069 watts with 12 h mode of day and night change.

### Bacteria isolation

In this study, we used strain *Janthinobacterium* sp. SLB01 was isolated from a sample of the diseased sponge *L. baicalensis*, collected in Lake Baikal, Central Siberia, Russia (Petrushin et al., 2019) for subsequent experimental infection, isolation, and identification using cell culture of the primmorphs. Healthy cell culture of the primmorphs (2–4 mm in diameter) of green color were transferred to the 24-well plates (Nalge Nunc International, Rochester, NY, USA), 1 pieces per well. The primmorphs were infected with 50 μl of *Janthinobacterium* sp. strain SLB01 in 2 ml of NBW (Natural Baikal drinking water). Infected the primmorphs were cultivated at 3–6°C with a 12-hour day and night cycle for 14 days. The cell suspension of the bacteria from infected primmorphs homogenized and passed through a MF-Millipore membrane filter, 0.45 μm pore size (Merck, Switzerland). The bacteria was cultured on nutrient media with R2A (0.05% yeast extract, 0.05% tryptone, 0.05% casamino acids, 0.05% dextrose, 0.05% soluble starch, 0.03% sodium pyruvate, 1.7 mM K2HPO4, 0.2 mM MgSO4, final pH 7.2 adjusted with crystalline K2HPO4 or KH2PO4) agar plates (Merck KGaA, Darmstadt, Germany) at 21°C, at pH 7.2. Bacterial cell morphology was determined using light microscopy Axio Imager Z2 microscope (Zeiss, Oberkochen, Germany) equipped with fluorescence optics (self-regulating, blue HBO 100 filter, 358/493 nm excitation, 463/520 nm emission). The isolates of bacteria were stained with a NucBlue Live ReadyProbes reagent (Thermo Fisher Scientific Inc., Waltham, MA, USA). During the infection, the observations were carried out with daily descriptions, as well as sampling for DNA isolation and sequencing.

### Microscopy studies, light and electron microscopy

We observed changes in infected cell cultures of primmorphs *Janthinobacterium* sp. strain SLB01 every day for 14 days. The samples of cell cultures were stained with a NucBlue Live ReadyProbes reagent (Invitrogen, USA). Cell morphology was determined by light microscopy Axio Imager Z2 microscope (Zeiss, Oberkochen, Germany) equipped with fluorescence optics (self-regulating, blue HBO 100 filter, 358/493 nm excitation, 463/520 nm emission). The strains of bacteria were stained with a NucBlue Live ReadyProbes reagent (Thermo Fisher Scientific Inc., Waltham, MA, USA).

The samples were prepared for scanning electron microscopy (SEM) analyses. Fixation was performed according to the following procedure: pre-fixation in 1% OsO4 (10 min), washing in cacodylate buffer (30 mM, pH 7.9) (10 min), fixation in 1.5% glutaraldehyde solution on cacodylate buffer (30 mM, pH 7.9) (1 h), washing in cacodylate buffer (30 mM, pH 7.9) (30 min), postfixation in 1% OsO4 solution on cacodylate buffer (30 mM, pH 7.9) (2 h), washing in filtered Baikal water 15 min at room temperature, and dehydration in a graded ethanol series. The specimens were placed into SEM stubs, dried to a critical point, and coated with liquid carbon dioxide (BalTec CPD 030) using a Cressington 308 UHR sputter coater before examination under Sigma series scanning electron microscope (Zeiss, Oberkochen, Germany) operated at 5.0 190 kV. The samples for transmission electron microscope (TEM) were taken (strain *Jantinobacterium* sp. bacteria, healthy primmorphs, primmorphs 24 hours, 3 and 7 days after infection with strain of the *Jantinobacterium* sp. were fixed with 2.5% glutaraldehyde in 0.1 M cacodillate buffer (pH 7.2) for 24 hours at 4°C. Then the material was washed in 0.1 M cacodillate buffer (pH 7.2) – 3 times for 1 hour, 1% OsO4 diluted in 0.1 M cacodillate buffer (pH 7.2) was fixed for 30 minutes. After washing from the fixative in distilled water (3 times for 30 minutes), the material was dehydrated in a series of ethanol of increasing concentration and acetone; then the material was enclosed in a mixture of Epon and Araldite (Sigma, USA). Semi-thin and ultra-thin sections were made using the Leica UC7 ultramicrotome (Leica Microsystems, Germany). Ultrathin sections were contrasted with a 0.5% aqueous solution of uranyl acetate (20 min.) and Reynolds lead citrate (10 min.). Ultrathin sections were analyzed using a Libra 200 FE transmission electron microscope (Carl Zeiss, Oberkochen, Germany) and Libra 120 (Carl Zeiss, Oberkochen, Germany).

### Genome sequencing

The DNA was isolated for full-genome sequencing and was performed the identity of the isolated strain. Genomic DNA was isolated following a standard bacterial DNA isolation cetyltrimethylammonium bromide (CTAB) protocol (https://jgi.doe.gov/user-programs/pmo-overview/protocols-sample-preparation-information/jgi-bacterial-dna-isolation-ctab-protocol2012/). The sequence library was prepared with the Nextera XT DNA library preparation kit (Illumina, USA). Genomes were sequenced on the Illumina MiSeq platform using v2 paired-end chemistry (2 × 250 bp, 12,099,942 reads total). The bacterial genus of the isolate was identified by analysis of sequencing data of the 400 universal marker genes.

### Genome assembly, annotation and phylogenetic relationship

Raw reads error correction and filtering with FastP tool with default settings (Chen et al., 2018). Genomes were assembled with SPAdes version 3.11.0 (Nurk et al., 2013) with default settings. Contigs from draft assembly with a length of more than 10 Kbp were scaffolded with Ragout version 2.2 with default settings (Kolmogorov et al., 2018), (https://github.com/fenderglass/Ragout) using *Janthinobacterium* sp. LM6 chromosome (GenBank accession no. CP019510) as the reference. Gene annotations were performed using PGAP (https://github.com/ncbi/pgap). Core genome construction was made with Roary version 3.13.0 with default settings (Page et al., 2015). Genome completeness analysis made with benchmarking universal single-copy orthologs (BUSCO) version 5.0.0 using dataset “burkholderiales_odb10” (Waterhouse et al., 2018). A phylogenetic tree for set of closer strains was constructed using PhyloPhlAn based on concatenated alignments of up to 400 conserved proteins (Asnicar et al., 2020) using “supermatrix_aa” and “low diversity” mode with the “phylophlan” database. Other settings were set to defaults.

## RESULTS

### Light and electron microscopy

We used a model cell culture of healthy primmorphs of sponge *L. baicalensis* for experimental infection with the *Janthinobacterium* sp. strain SLB01 isolated from a diseased sponge the *L. baicalensis* to determine this strain’s pathogenicity. The healthy primmorphs are bright green in color and bright red autofluorescence of chlorophyll in the cells due to green symbiotic microalgae belonging to the taxon Chlorophyta in their composition. We observed cells of the sponges, a strict arrangement of symbiotic microalgae in the amoebocytes with nuclei in uninfected primmorphs (**Figures 1A,B**). A completely different picture was observed in infected primmorphs with *Janthinobacterium* sp. strain SLB01. Dirty scurf, fetid odor, and biofilm formation were observed in the infected cultures, which was probably associated with the growth of bacteria. All samples of primmorphs lost their green color; there was a chaotic arrangement and adhesion of microalgae, an imbalance in cells of sponges, observed were destroyed of amoebocytes after being infected with the strain *Janthinobacterium* sp. SLB01 (**Figure 1C**). We observed by fluorescence microscopy suppression of the autofluorescence chlorophyll-containing intracellular of microalgae in infected primmorphs with *Janthinobacterium* sp. strain SLB01 on day 3 of cultivation (**Figure 1D**). The dead cells of sponges and chaotic arrangement of microalgae were observed with further cultivation of the infected culture of the primmorphs with an increase in the number of short rod-shaped bacteria after 7 days of cultivation (**Figures 1E,F**). We found that short rod-shaped bacteria surrounded cells of microalgae. The loss of chlorophyll autofluorescence and the death of microalgae were observed in all the experimental samples.

**Figure 1.**
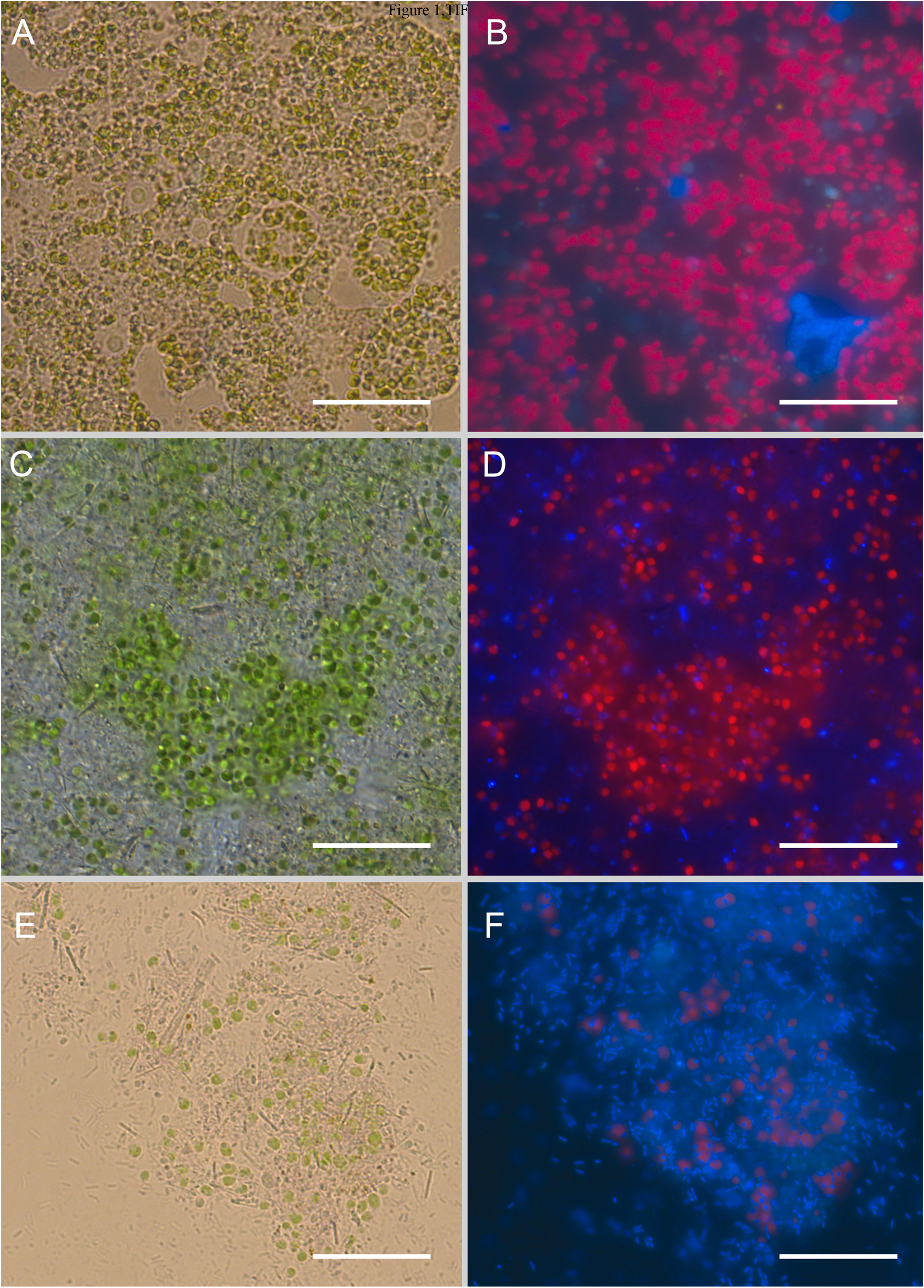
Light and fluorescence images of cell culture of the primmorphs of sponge *L. baicalensis*. (A) Green microalgae are located within amoebocytes in the healthy cell culture of primmorphs. (B) Fluorescence microscopy shows red autofluorescence of chlorophyll-containing intracellular microalgae in the healthy primmorphs. (C) The death of green microalgae and the increase of bacteria on day 3 of cultivated are shown in the primmorphs infected with *Janthinobacterium* sp. strain SLB01 (indicated by arrow). (D) Fluorescence microscopy shows the death of microalgae (red color) in infected primmorphs with *Janthinobacterium* sp. strain SLB01 on day 3. Bacteria are shown with blue color. (E) The primmorphs infected with *Janthinobacterium* sp. strain SLB01 during 7 days, death of green symbionts observed and huge biomass of bacteria (shown by arrows). (F) Fluorescence microscopy shows the death of microalgae in the primmorphs and increases bacterial biomass. Bacteria are shown with blue color. The samples of primmorphs were stained with the NucBlue Live ReadyProbes reagent for fluorescence microscopy. Scale bars: 10 μm.

We observed the interaction of bacteria with the host cells when using SEM microscopy in the infected primmorphs with *Janthinobacterium* sp. strain SLB01 (**Figure 2B**). The squamous epithelium was destroyed; the symbiotic microalgae were packed entirely in a thick microbial biofilm with short rod-shaped bacteria (**Figure 2B**). In addition, when cultivating strain *Janthinobacterium* sp. SLB01, we experimentally observed biofilms and floc formation in the infected cell cultures of primmorphs of *L. baicalensis*. While, in the healthy primmorphs, the epithelium surface was clean, even, and smooth, the microalgae contained were spheroidal and 2.5-3.0 μm in diameter with a clean cell wall (**Figure 2A**).

**Figure 2.**
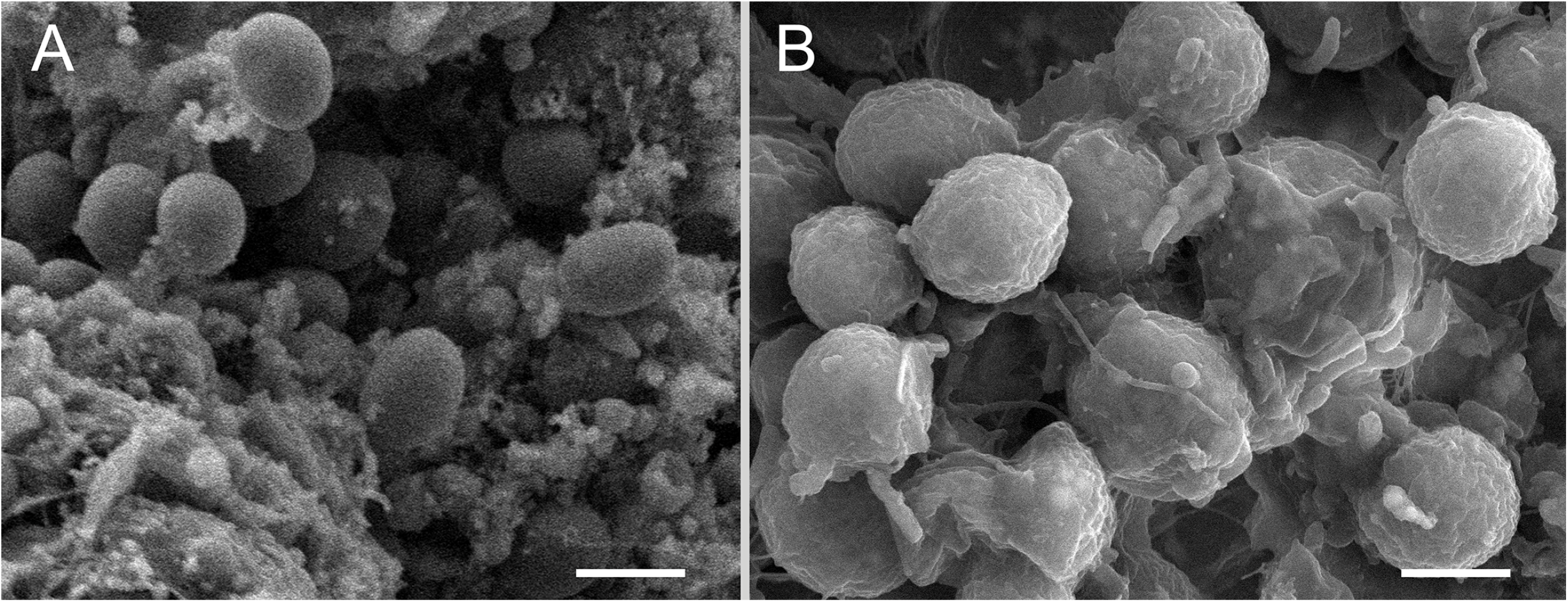
SEM images of cell cultures of the primmorphs. (a) The healthy primmorphs not infected with *Janthinobacterium* sp. strain SLB01 (shown by arrows). (b) The primmorphs infected with *Janthinobacterium* sp. strain SLB01. The formation of microbial biofilm with short rod-shaped bacteria on day 7th of cultivation on the surface of symbiotic microalgae was observed. Scale bars are 2 μm.

In more detail, we observed the infection course when using ultrastructural analysis of the TEM microscopy of the interaction of bacteria *Janthinobacterium* sp. strain SLB01 with cell cultures of primmorphs of the sponge *L. baicalensis* (**Figure 3**). We found that amoebocyte cells filled with green symbiotic microalgae in cultures of healthy primmorphs (**Figure 3A**). These are cells up to 20 μm in diameter containing a nucleus with a prominent nucleolus. The cytoplasm of amoebocytes contains dictyosomes of the Golgi apparatus and cisterns of the endoplasmic reticulum. A distinctive feature of amoebocytes is the presence in the cytoplasm of specialized vacuoles - symbiosomes with symbiotic microalgae representatives of the *Chlorophyceae* family enclosed in them. Microalgae cells with a diameter of 2.5-3.0 μm have a thin electron-dense polysaccharide envelope separated from the cell’s outer membrane by a narrow supramembrane space (**Figure 3B**). There is a chloroplast, which may contain electron-transparent inclusions; obviously, starch grains. Storage starch granules are often present between the thylakoid membranes. These granules are often present between the thylakoid membranes and directly in the cytoplasm of microalgae (**Figure 3B**). In addition, it was noted that no found bacteria in the mesoglea of healthy primmorphs. The rod-shaped bacteria were found in mesoglea of cell cultures of the primmorphs 24 hours after infection (**Figure 3D**). The structure of bacteria changes in comparison and they had an enlarged folded outer membrane. There was an electron-transparent hallo around the bacterial cells, which indicates their ability to lyse the surrounding mesoglea components. In addition to mesoglea, bacteria are present in amoebocytes, where they penetrate by phagocytosis. The destruction of the system of intracellular membranes was observed and vacuolization enhanced. At this stage of the experiment, symbiosomes with microalgae enclosed in them were preserved in the cytoplasm of amoebocytes. However, some symbiotic microalgae leave the host cells and are located in the mesoglea. The destruction processes of primmorphs cells reach the terminal stage on the 7th day of infection (**Figure 3E**). The 7 days after the start of the infection, microalgae were located directly in the mesoglea, where they become available for the action of bacteria. The cytoplasm of primmorphs was fragmented, resulting in the absence of whole, functionally active cells. We observed that the mesoglea also contains symbiotic microalgae infected with bacteria, which then, due to division, form colonies of microorganisms united by contact processes (**Figure 3F**). The formation of bacterial colonies was accompanied by the utilization of the components of the microalgae cytoplasm and there was only a polysaccharide shell with the bacteria enclosed in it (**Figure 3G**). Thus, as the infection progresses, the primmorphs cells lyse and the microalgae are localized mainly in the mesoglea. During this period, they become infected with bacteria, which eventually leads to the formation of colonies of microorganisms inside the microalgae shell on the 7th day of infection.

**Figure 3.**
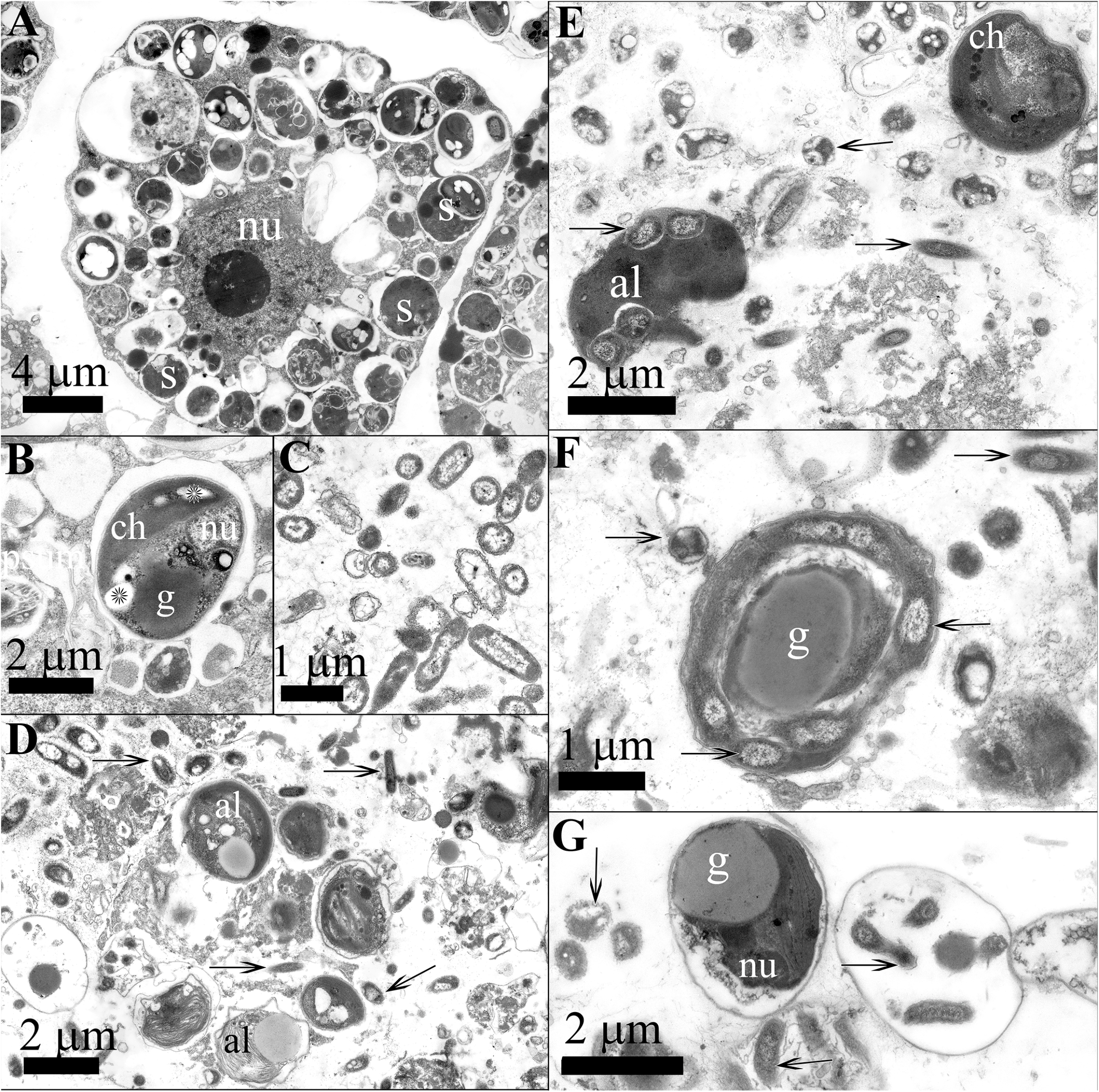
Ultrastructural analysis of *Janthinobacterium* sp. strain SLB01 with cell culture of primmorphs of the sponge *L. baicalensis*. (a) The amebocyte of healthy primmorph with symbiotic microalgae in the cytoplasm. (b) Symbiosome from amebocyte of healthy primmorph with the microalgae contained a nucleus, chloroplast, and inclusions, including osmiophilic lipid granules and starch grains (marked with asterisks). (c) The *Janthinobacterium* sp. strain SLB01 bacterial culture was used in experiment. (d) The structure of the primmorphs of the sponge *L. baicalensis*, one day after infection, the presence of Gram-negative bacteria (marked with black arrows) in the mesoglea and amebocytes. (e-f) The development of the infectious process on the 7th day of the experiment. (g) Host cells undergo lysis, and bacteria penetrate free-lying microalgae and form colonies. Legend: al - microalgae cell; ch - chloroplast; l - primary lysosome, lg - lipid granule; nu - nucleus; ph – phagosome

### Isolation strain of *Janthinobacterium* sp. from infected primmorphs

We isolated bacteria from the cell culture of the primmorphs infected by strain *Janthinobacterium* sp. SLB01. That was rod-shaped, motile, aerobic bacteria; a purple pigment violacein appeared on the second day. A morphological analysis of the strain showed that bacteria have short-rods up to 2.0 μm long and 0.3 μm in diameter, with a two-layer outer membrane typical of Gram-negative bacteria (**Figure 3C**). The cytoplasm, in most cases, is granular, electron-dense, the nucleoid zone is well defined, and there were flagella.

### Comparison genomes of strains *Janthinobacterium* sp

Genome assembly and annotation with NCBI Prokaryotic Genome Annotation Pipeline (PGAP) for *Janthinobacterium* sp. PLB02 was done as described previously (Petrushin et al., 2017). The strain *Janthinobacterium* sp. PLB02 was isolated from infected primmorphs with the *Janthinobacterium* sp. strain SLB01. In addition, we carried out sequencing and analysis of the genomes of *Janthinobacterium* sp. strain SLB01 and new strain *Janthinobacterium* sp. PLB02 from infected primmorphs. We assembled the genome of *Janthinobacterium* sp. strain PLB02 from raw data the same way as for strain *Janthinobacterium* sp. SLB01. Genomes were released in NCBI for further study and annotation. In the present study, we compared the genomic content of two strains *Janthinobacterium* sp. SLB01 and isolated *Janthinobacterium* sp. PLB02. We used the same software set for genome assembly and annotation to prevent genome variations depending on reference and assembly software versions. Although the PGAP version for annotation *Janthinobacterium* sp. SLB01 was 4.13 (used when genome released in NCBI RefSeq). To annotate *Janthinobacterium* sp. PLB02, we used the current available NCBI PGAP version 5.1, which can annotate more genes because its database is richer. The final genome assembly statistics of raw reads count, genome size, number of genes, pseudogenes, protein-coding sequences, tRNA noncoding RNA, and references to genome reports are presented (**Table 1**). Genome completeness analysis with benchmarking universal single-copy orthologs (BUSCO) (Waterhouse et al., 2018) showed results for both: *Janthinobacterium* sp. strain SLB01 and strain PLB02 99.1 % complete, no fragmented, and 0.9% missing BUSCOs. The fully assembled genomes comprised 6,467,981bp for strain *Janthinobacterium* sp. SLB01 and 6,417,505 bp for strain *Janthinobacterium* sp. PLB02 and exhibited similar G+C content (62.63% and 62.65%, respectively). Genome annotation with PGAP revealed 5,612 genes (5,498 protein-coding) for strain SLB01 and 5,632 (5,518 protein-coding) for strain PLB02, respectively in Table 1. We compared the genomic content of the *Janthinobacterium* sp. strain SLB01 and *Janthinobacterium* sp. strain PLB02 with Roary (Page et al., 2015) and revealed most of the genes are the same (have a homology of more than 99%). We built a phylogenetic tree for some of these species to compare genomic features of both strains *Janthinobacterium* sp. SLB01 and PLB02 with closer species (Haack et al., 2016). Strain *Janthinobacterium* sp. PLB02 showed the highest 400 genes phylogenetic affiliation to strain *Janthinobacterium* sp. SLB01 of phylum *Proteobacterium*, *Betaproteobacteria* of the family *Oxalobacteraceae* (**Figure 4**). Bacterial strains of *Janthinobacterium* sp. SLB01 and *Janthinobacterium* sp. PLB02, based on a comparison of complete genomes, showed 99 % similarity with strain *J. lividum* MTR.

**Figure 4.**
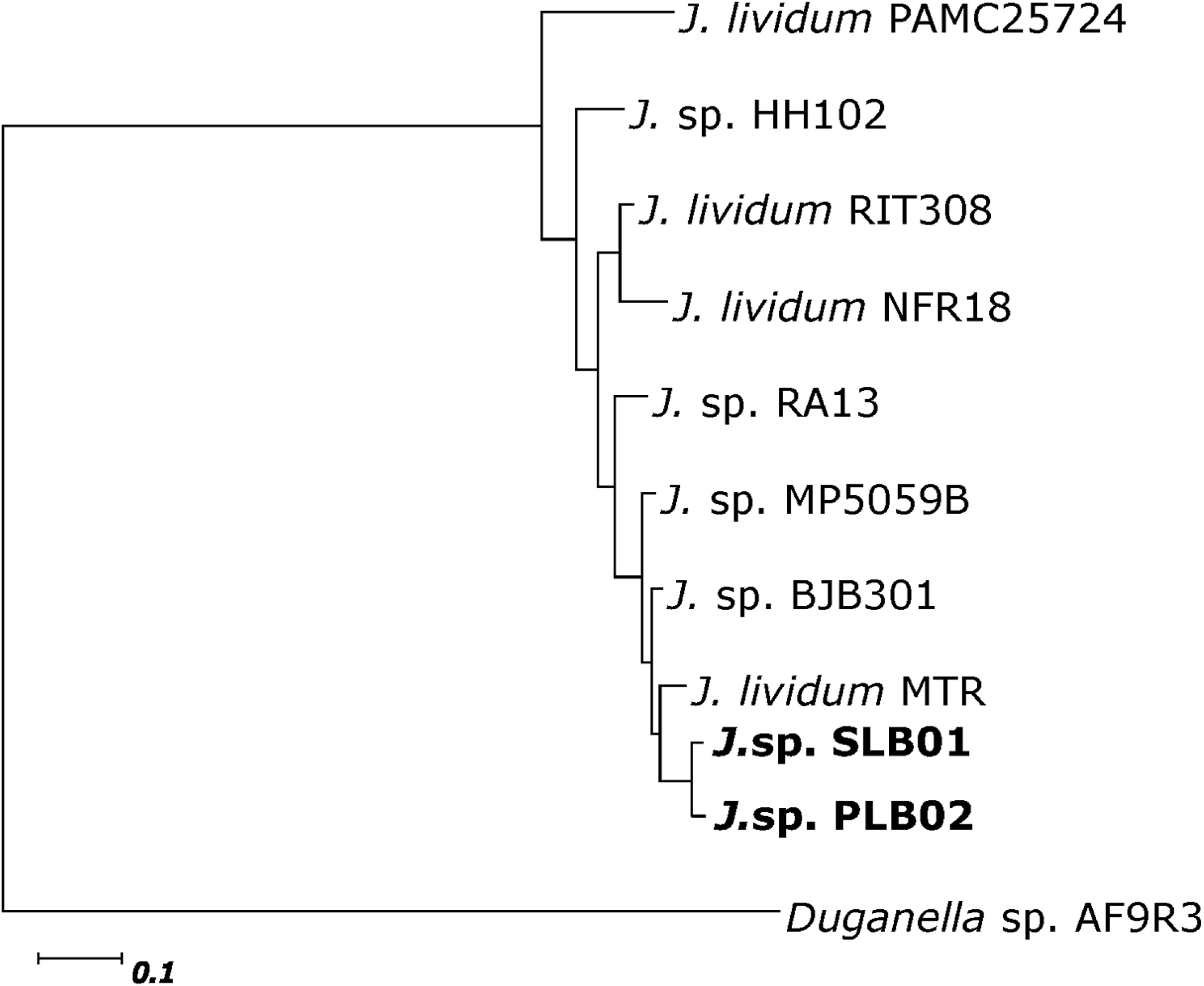
Phylogenetic tree of strains *Janthinobacterium* sp. SLB01 and *Janthinobacterium* sp. PLB02 with closely related species. Trees are built based on approximately 400 universal marker genes by PhyloPhlAn (a maximum-likelihood method). The genome properties of each strain in this study are presented in **Table 1**.

We compared the obtained genomes of strains of the *Janthinobacterium* sp. PLB02 and *Janthinobacterium* sp. SLB01 with each other, analyzed coding virulence proteins, and key genes and such as genes of violacein, floc formation, and the type VI secretion system. We found, that these strains are indistinguishable and 100% homologous to each other in these virulence factors. We showed earlier that the strain *Janthinobacterium* sp. SLB01 was able to produce violacein and contain violacein synthesis operon vioABCDE (Petrushin et al., 2020). Isolated from primmorphs, the strain *Janthinobacterium* sp. PLB02 also produces a pigment violacein, and the genome contains violacein synthesis operon vioABCDE. The gene coordinates and locus names flanking regions are presented (**Figure 5** and Table S1). We found 100% homology of these strains except for the vioD gene, since the reading of the genome of the strain *Janthinobacterium* sp. PLB02 began with second methionine. An annotated name for the genes of violacein synthesis operons was discovered, instead of the generally accepted names for violacein synthesis operon vioABCDE at comparing their genomes (Table S1).

**Figure 5.**
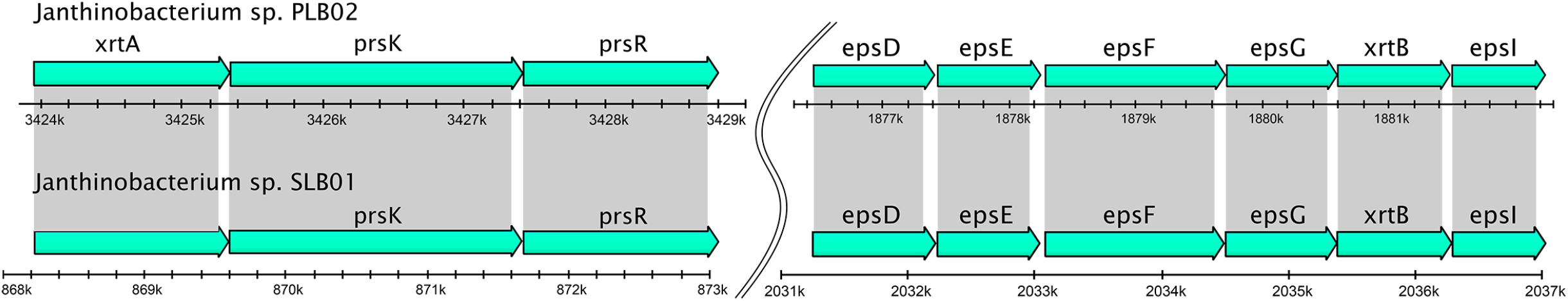
Comparison diagram of the violacein production loci and flanking regions in the *Janthinobacterium* sp. PLB02 (loci prefix J3P46) and *Janthinobacterium* sp. SLB01 (loci prefix F3B38) genomes. Genes are displayed with arrows, loci tags signed with regular and gene names signed with an italic font. Violacein operons are displayed with the semitransparent violet bar.

The previous study discovered gene clusters involved in floc formation (Petrushin et al., 2020). We examined whether the isolated *Janthinobacterium* strain PLB02 had genes of the floc formation. It was found, that these clusters of genes strain the *Janthinobacterium* sp. PLB02 has an identity of 100% structure genome to the *Janthinobacterium* sp. strain SLB01 (**Figure 6**). Earlier, we showed that *Janthinobacterium* sp. SLB01 formed a strong biofilm that was rich in exopolysaccharides (EPS) in the stationary phase. It was noted that the isolated strain *Janthinobacterium* sp. PLB02 also formed a strong biofilm. This strain produces floc and strong biofilm by exopolysaccharide biosynthesis and PEP-CTERM/XrtA protein expression. Genome analysis showed that strain *Janthinobacterium* sp. PLB02 revealed all of the required gene clusters for floc formation. Its genome has system glycosyltransferase putative exosortase XrtA (previously called EpsH), PEP-CTERM system histidine kinase PrsK, PEP-CTERM system associated sugar transferase, sensor histidine kinase of a two-component system and PEP-CTERM-box response regulator transcription factor PrsR. Localization, annotation, and identity percentage of these genes are presented (Table S2). In addition, it was shown, that the genome of *Janthinobacterium* sp. SLB01 contains all three categories of the genes required to function of the type VI secretion system (T6SS) as the primary virulence factor. We analyzed the type VI secretion system in the genome of the strain *Janthinobacterium* sp. PLB02 and found that the genes of both strains are 100% identical to each other. The genome of the isolated strain the *Janthinobacterium* sp. PLB02 also contains all three categories of the genes required for the function of the type VI secretion system. These genes *Janthinobacterium* sp. strain PLB02 are allocated through the genome by 10 clusters, as in the strain *Janthinobacterium* sp. SLB01. Thus, the isolated strain *Janthinobacterium* sp. PLB02 was identical to strain *Janthinobacterium* sp. SLB01, and these strains are pathogens for *L. baicalensis* sponges and cell culture of primmorphs.

**Figure 6.**
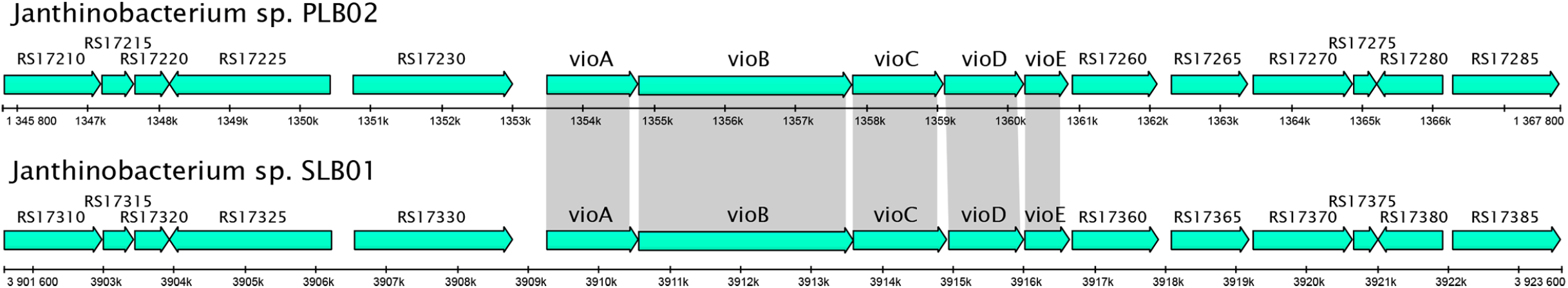
Schematic diagram of the genetic organization of *Janthinobacterium* sp. strain PLB02 and *Janthinobacterium* sp. strain SLB01 gene clusters required for floc formation: for exopolysaccharides (EPS) synthesis, PEP-CTERM, and exosortase. Arrows indicate genes and the direction of the arrows represents the direction of transcription of the genes in the genome, gene names signed with an italic font.

## DISCUSSION

Here we have shown that the strain Janthinobacterium sp. SLB01 isolated from the diseased sponge and infected cell culture of the primmorphs is the same and the genomes of these strains are almost identical. Strains of *Janthinobacterium* sp. SLB01 and *Janthinobacterium* sp. PLB02 are pathogens for cell cultures of primmorphs and *L. baicalensis* sponges. We found at the experimental infection of cell culture of the primmorphs that short rod-shaped bacteria of the strain *Janthinobacterium* sp. SLB01 grown quickly and are parasitizes both sponge cells and symbiotic microalgae. We detected a mass death of the symbiont microalgae (Chlorophyta) and sponge cells in the infected primmorphs and increased bacteria of the strain the *Janthinobacterium* sp.

The bacteria *Janthinobacterium* sp. was founded in mesoglea of cell cultures of the primmorphs from 24 hours after infection and were able to lyse the cells of the primmorphs. The characteristic features of the structure of *Janthinobacterium* sp. during the development of the infectious process were the presence of a folding outer membrane, an increase of the periplasm, and an electron-transparent zone of lysis around bacterial cells. It is known, that the outer cell membrane and the periplasm of Gram-negative bacteria is a compartment responsible for the production of secondary metabolites, including proteolytic enzymes and other factors of bacterial cell virulence (Teeseling et al., 2017; Miller and Salama, 2018). The increase in the surface area of the outer membrane and the volume of the periplasm in *Janthinobacterium* sp. in the course of infection indicates the activation of processes aimed at realizing their pathogenic potential. The *Janthinobacterium* sp. penetrate the cytoplasm of microalgae, and utilize their contents, using nutrients for growth, division, and formation of colonies of the bacteria. Thus, as the infection progresses in the cells of the primmorphs and the microalgae the processes of reaching the terminal stage on the 7 days of infection indicate a rather aggressive state of bacteria the rapid course of the pathogenic process. We observed the destruction of the photosynthetic apparatus, the loss of chlorophyll autofluorescence, and the death of symbiotic microalgae in all the infected primmorphs.

We have previously shown that there is a significant increase in the number of opportunistic microorganisms, including *Betaproteobacteria* of the *Oxalobacteraceae* family in diseased freshwater sponges was detected by analysis of 16S rRNA genes (Belikov et al., 2019; Chernogor et al., 2020). We isolated, sequenced, and annotated the genome of the strain *Janthinobacterium* sp. SLB01 from diseased sponge collected from the Lake Baikal as previously described (Petrushin et al., 2019; 2020). It has been shown that the genome of this strain contains virulence factors. Therefore, it was decided to test the effect of this strain at the cell culture of the primmorphs, followed by isolation, sequencing, and analysis of the isolated genome of the *Janthinobacterium* sp. strain PLB02. Earlier, we showed that the cell culture of primmorphs of a healthy sponge *L. baicalensis* is identical to sponges and is a good cell culture as a model system for studying the disease of Baikal sponges (Chernogor et al., 2020). Developing a model to investigate the transmission of pathogenic agents from diseased sponges requires a detailed study of pathogen-host interactions in the environment. However, these experiments with sponges are difficult to carry out in the natural conditions of Lake Baikal, and therefore the primmorphs cell culture was used.

Comparing the two genomes of strains the *Janthinobacterium* sp. SLB01 and *Janthinobacterium* sp. PLB02 isolated from the diseased sponge and infected cell culture of the primmorphs with the same strain shows that genomes of these bacteria have identical genomic content. The genome size, genes count, and G+C content is very close. The genome size of *Janthinobacterium* strains slightly differs because of the number of Ns (unknown nucleotides) after the scaffolding procedure. Core genome construction shows that most genes (5485 of 5557) are identical about 99 %. A similar strain *Janthinobacterium* sp. PLB02 was isolated and identified from the infected cell culture of the primmorphs. It is known that these are rod-shaped Gram-negative bacteria that produce violacein, a compound with antimicrobial and antiviral properties that is toxic to eukaryotic cells (Matz et al, 2008). The isolated bacteria *Janthinobacterium* sp. PLB02 can colonize the space and possibly suppress the grown microalgae with pigment violacein. This pigment production was observed in infected primmorphs and all the genes (operon VioABCDE) are present in its genome. We identified five genes encoding VioA, VioB, VioC, VioD, and VioE proteins related to violacein biosynthesis like those identified in published *Janthinobacterium* sp. SLB01. Earlier observed that one essential strategy of bacteria *Jantinobacterium* sp. strain SLB01 is the secretion of virulence factors through the cell membranes of the victim to achieve a potential target (Petrushin et al., 2020). Here we showed that the bacteria attacked eukaryotic cells of the microalgae, and then took released nutrients after cell lysis during the experimental infection of the cell culture of primmorphs with bacterial strain *Janthinobacterium* sp. SLB01 (**Figures 3F,G**). In addition, an identical T6SS secretion system of the strain *Jantinobacterium* sp. SLB01 was found in the isolated *Janthinobacterium* sp. PLB02. Both strain’s genomes contain all three categories of genes required for the function of the type VI secretion system (T6SS) (Silverman et al., 2012; Wang et al., 2019). Therefore, their so fast and aggressive state is explained in the destruction of host cells of the primmorphs.

The result of the phylogenetic tree based on 400 universal marker genes by PhyloPhlAn (a maximum-likelihood method) shown that the genomes of *Janthinobacterium* sp. SLB01 and *Janthinobacterium* sp. PLB02 is homologous to each other and the closest related to the psychrotolerant strain *Janthinobacterium lividum* MTR (**Figure 4**). Interesting, that *J. lividum* either cause necrosis on mushroom tissue blocks and colonizes the skin of some amphibians confers protection against fungal pathogens (Gillis and Logan, 2015; Valdes et al., 2015). In addition, isolated bacteria can also produce floc formation and strong biofilm in the stationary phase. When cultivating the strains *Janthinobacterium* sp. SLB01 and *Janthinobacterium* sp. PLB02, we observed biofilms and floc formation in the diseased sponges and the infected cell cultures of the primmorphs of *L. baicalensis*. When genomic analysis of the two strains, we found RpoN, PepA, XrtA, PrsK, PrsR gene clusters for the formation of floc and 100% their identity in the strains (Table S2). Using ultrastructural analysis, we found that the symbiotic microalgae were completely packed in a thick microbial biofilm at infected of the primmorphs the strain *Jantinobacterium* sp. SLB01 (**Figure 2B**). Moreover, on the 7th day of infection, we discovered the formation of bacterial colonies was accompanied by the utilization of the components of the microalgae cytoplasm and there was only a polysaccharide shell with the bacteria enclosed in it (**Figure 3G**). This proves how fast and aggressive the action of these bacteria is. Thus, floc formation and biofilm can negatively affect the physiology of life of the host (sponge *L. baicalensis*) due to clogging of the pores. Earlier these negative effects of biofouling on the functioning of the filter-feeding sponge *Halisarca caerulea* has been shown (Alexander et al., 2015). The Exopolysaccharides (EPS) are known to be the main component of biofilm produced by the species of the *Oxalobacteraceae* (Pantanella et al., 2007).

It is known that bacteria of the family *Oxalobacteraceae* is characterized by the presence of ecologically extremely diverse organisms and contains environmental saprophytic organisms, phytopathogens, and opportunistic pathogens, including those common for freshwater ecosystems (Coenye, 2014). The genomes of many environmental isolates of *Janthinobacterium* from ice, waters, sediments, and soils were sequenced (Garrity et al., 2005; Gillis and De Ley, 2006; Gillis and Logan, 2015), but strains of *Janthinobacterium* sp. strain SLB01 and new *Janthinobacterium* sp. strain PLB02 from the Baikal sponge and primmorphs were isolated for the first time.

Disease and mass mortality of sponges and corals have been observed worldwide in the marine environment in recent years (Webster et al., 2004; Olson et al., 2006; Hoegh-Guldberg et al., 2007; Miller, and Richardson, 2011; Pita et al., 2018). The die-off events threaten the entire sponge-associated biodiversity (Olson et al., 2006; Webster, 2007; Stabili et al., 2012). These changes in sponge-microbe interactions appear associated with climate change and the occurrence of opportunistic infections resulting from changes in water temperature caused by global warming, light intensity, and salinity (Webster et al., 2008; Luter et al., 2010; Erwin et al., 2012; Fan et al., 2013; Fujise et al., 2014). Previously, was shown isolation and description of the pathogenic bacterial strain NW4327 from an infected marine sponge *Rhopaloeides odorabile* in the Great Barrier Reef (Webster et al., 2002). Researchers reported the isolation of the pathogenic bacterial strain of *Pseudoalteromonas agarivorans* that has been found in diseased sea sponges and has pathogenicity genes (Choudhury et al., 2015).

Thus, in this study, we tried to reproduce Koch’s postulates with the cell culture of the primmorphs. The present study is the first; we were able to isolate a new strain Janthinobacterium sp. PLB02, after infecting cell culture of the primmorphs of sponge L. baicalensis using the original strain Janthinobacterium sp. SLB01. The strain Janthinobacterium sp. SLB01 was isolated from diseased sponges. These strains are practically the same and have virulence factors in their genomes. In more detail, we showed this bacterium is a potential pathogen for Baikal sponges and primmorphs. The study results will help to expand the understanding of microbial relationships in the development of disease and the death of Baikal sponges.

## Contribution to the field

1- Expand understanding about symbiotic relationships in the freshwater sponges. 2- Allow increase understanding of the pathogenicity of Janthinobacterium sp. strains for sponges. 3- Janthinobacterium sp. produces pigment violacein, what synthesis operon vioABCDE. 4- Strains Janthinobacterium sp. SLB01 and PLB02 produce floc formation and strong biofilm. 5- Both strains genes are 100% identical and have the type VI secretion system in the genomes.

## Ethics statements

### Studies involving animal subjects

Generated Statement: No animal studies are presented in this manuscript.

### Studies involving human subjects

Generated Statement: No human studies are presented in this manuscript.

### Inclusion of identifiable human data

Generated Statement: No potentially identifiable human images or data is presented in this study.

## Data availability statement

Generated Statement: The datasets presented in this study can be found in online repositories. The names of the repository/repositories and accession number(s) can be found below: https://www.ncbi.nlm.nih.gov/, NZ_CP071520.

## AUTHOR CONTRIBUTION

Conceptualization, S.B. and L.C.; methodology, L.C. and M.E., software, I.P. formal analysis, L.C., S.B., I.P.; investigation, L.C., S.B., M.E., I.P., and I.K.; data curation, I.P.; writing—Original draft preparation, L.C., M.E., S.B., I.P., E.C., and writing— Review and editing, L.C., S.B., M.E., E.C. and I.P.; visualization, L.C., M. E., I.P.; supervision, S.B. and L.C.; funding acquisition, S.B. All authors have read and agreed to the published version of the manuscript.

## ACKNOWLEDGMENTS

This research and article processing charge was funded by the Russian Science Foundation, grant number 19-14-00088. Sample collection of sponges was carried out within the framework of the Siberian Branch, Russian Academy of Sciences basic budget funding from No 0345-2019-0002. Complete sequencing of bacterial genomes was performed in the Center of Shared Scientific Equipment “Persistence of microorganisms” of ICIS UB RAS, Russia. The authors appreciate the help of the service staff of the National Scientific Center of Marine Biology FEB RAS (the former AV Zhirmunsky Institute of Marine Biology).

## Data Availability

This full-genome shotgun project has been deposited at DDBJ/ENA/GenBank and in the Sequence Read Archive (SRA) and published with accession number NZ_CP071520.

## Abbreviations

BUSCO: Benchmarking universal single-copy orthologs
CTERM: C-terminal
EPS: Exopolysaccharides
PGAP: Prokaryotic genome annotation pipeline
PEP: Pro-Glu-Pro
PUL: Polysaccharides utilization loci
T6SS: Type VI secretion system
Floc: Floc formation

## References

Alexander, B. E., Mueller, B., Vermeij, M. J. A., van der Geest, H. H. G., and de Goeij, J. M. (2015). Biofouling of inlet pipes affects water quality in running seawater aquaria and compromises sponge cell proliferation. Peer J. 3, e1430. doi: 10.7717/peerj.1430

Asnicar, F., Thomas, A. M., Beghini, F., Mengoni, C., Manara, S., Manghi, P., et al. Precise phylogenetic analysis of microbial isolates and genomes from metagenomes using PhyloPhlAn 3.0. Nature Communications 2020, 11. doi: 10.1038/s41467-020-16366-7

Belikov, S., Belkova, N., Butina, T., Chernogor, L., Kley, A. M., Van; Nalian, A., et al. (2019). Diversity and shifts of the bacterial community associated with Baikal sponge mass mortalities. PLoS One 14, e0213926. doi: 10.1371/journal.pone.0213926

Bil, K., Titlyanov, E., Berner, T., Fomina, I., and Muscatine, L. (1999) Some aspects of the physiology and biochemistry of *Lubomirska baicalensis*, a sponge from Lake Baikal containing symbiotic algae. Symbiosis 26, 179–191.

Bingle, L. E. H., Bailey, C. M., and Pallen, M. J. (2008). Type VI secretion: a beginner’s guide. Curr. Opin. Microbiol. 11, 3–8. doi: 10.1016/j.mib.2008.01.006

Chen, S., Zhou, Y., Chen Y., and Gu, J. (2018). Fastp: An ultra-fast all-in-one FASTQ preprocessor. Bioinformatics 34, i884–i890. doi: 10.1093/bioinformatics/bty560.

Chernogor, L. I., Denikina, N. N., Belikov, S. I., and Ereskovsky, A. V. (2011). Long-term cultivation of primmorphs from freshwater Baikal sponges *Lubomirskia baicalensis*. Marine Biotechnol. 13, 782–792. doi: 10.1007/s10126-010-9340-9

Chernogor, L., Denikina, N., Kondratov, I., Solovarov, I., Khanaev, I., Belikov, S., and Ehrlich, H. (2013). Isolation and identification of the microalgal symbiont from primmorphs of the endemic freshwater sponge *Lubomirskia baicalensis* (Lubomirskiidae, Porifera). Eur. J. Phycol. 48, 497–508. doi: 10.1080/09670262.2013.862306

Chernogor, L., Klimenko, E., Khanaev, I., and Belikov, S. (2020). Microbiome analysis of healthy and diseased sponges *Lubomirskia baicalensis* by using cell cultures of primmorphs. PeerJ. 8, e9080. doi: 10.7717/peerj.9080

Choudhury, J. D., Pramanik, A., Webster, N. S., Llewellyn, L. E., Gachhui, R., and Mukherjee, J. (2015). The pathogen of the Great Barrier Reef sponge *Rhopaloeides odorabile* is a new strain of *Pseudoalteromonas agarivorans* containing abundant and diverse virulence-related genes. Marine Biotechnol. 17, 463–478. doi: 10.1007/s10126-015-9627-y

Coenye T. (2014). “The family *Burkholderiaceae*,” in The Prokaryotes: Alphaproteobacteria and Betaproteobacteria, eds E. Rosenberg, E. F. DeLong, S. Lory, E. Stackebrandt, and F. Thompson (New York: Springer-Verlag.). 759–76. doi: 10.1007/978-3-642-30197-1_239 doi: 10.1007/s00442-007-0910-0.

Durán, N., Justo, G. Z., Ferreira, C. V., Melo, P. S., Cordi, L., and Martins, D. (2007). Violacein: properties and biological activities. Biotechnol. Appl. Biochem. 48, 127. doi: 10.1042/BA20070115

Erwin, P. M., Pita, L., López-Legentil, S., and Turon X. (2012). Stability of sponge-associated bacteria over large seasonal shifts in temperature and irradiance. Appl. Environ. Microbiol. 78, 7358–7368. doi: 10.1128/AEM.02035-12

Fan, L., Liu, M., Simister, R., Webster, N. S., and Thomas, T. (2013). Marine microbial symbiosis heats up: the phylogenetic and functional response of a sponge holobiont to thermal stress. The ISME Journal 7, 991–1002. doi: 10.1038/ismej.2012.165

Fieseler, L., Horn, M., Wagner, M., and Hentschel, U. (2004). Discovery of the novel candidate phylum ‘‘*Poribacteria*’ in marine sponges. J. Appl. Envion. Microbiol. 70, 3724–3732. doi: 10.1128/AEM.70.6.3724-3732.2004

Fujise, L., Yamashita, H., Suzuki, G., Sasaki, K., Liao, L. M., and Koike, K. (2014). Moderate thermal stress causes active and immediate expulsion of photosynthetically damaged Zooxanthellae (*Symbiodinium*) from corals. PLoS ONE 9, e114321. doi: 10.1371/journal.pone.0114321.g007

Gao, N., Xia, M., Dai, J., Yu, D., An, W., Li, S., et al. (2018). Both widespread PEPCTERM proteins and exopolysaccharides are required for floc formation of *Zoogloea resiniphila* and other activated sludge bacteria. Environ. Microbiol. 20, 1677–1692. doi: 10.1111/1462-2920.14080

Garrity, G. M., Bell, J. A., Lilburn, T. E. and Family, I. I. (2005). “*Oxalobacteraceae* fam. nov.,” in Bergey’s Manual of Systematic Bacteriology, eds G. M. Garrity, D. J. Brenner, N. R. Krieg and J. Staley (New York: Springer). 2, 623. doi: 10.1002/9781118960608.fbm00183

Gillis, M. and Logan, N. A. (2015). “Janthinobacterium,” in Bergey’s Manual of Systematics of Archaea and Bacteria, eds M.E. Trujillo, S. Dedysh, P. DeVos, B. Hedlund, P. Kämpfer, F.A. Rainey and W.B. Whitman. doi.org/10.1002/9781118960608.gbm00964

Gillis, M., and De Ley, J. (2006). “The Genera *Chromobacterium* and *Janthinobacterium*,” in The Prokaryotes: Handbook on the Biology of Bacteria, eds M. Dworkin, S. Falkow, E. Rosenberg, K. H. Schleifer and E. Stackebrandt (New York, Springer). 7, 737. doi: 10.1007/0-387-30745-1_32

Haack, F. S., Poehlein, A., Kröger, C., Voigt, C. A., Piepenbring, M., Bode, H. B., et al. (2016). Molecular keys to the *Janthinobacterium* and *Duganella spp*. interaction with the plant pathogen *Fusarium graminearum*. Front. Microbiol. 7. doi: 10.3389/fmicb.2016.01668

Hedges, S. B., Blair, J. E., Venturi, M. L., and Shoe, J. L. (2004). A molecular timescale of eukaryote evolution and the rise of complex multicellular life. BMC Evol. Biol. 4, 2. doi: 10.1186/1471-2148-4-2

Hentschel, U., Fieseler, L., Wehrl, M., Gernert, C., Steinert, M., Hacker, J., and Horn, M. (2003). “Microbial diversity of marine sponges,” in Molecular marine biology of sponges, ed W. E. G. Muller (Heidelberg, Germany: Springer-Verlag). 60–88. doi: 10.1007/978-3-642-55519-0_3

Hoegh-Guldberg, O., Mumby, P. J., Hooten, A. J., Steneck, R. S., Greenfield, P., Gomez E., et al. (2007). Coral reefs under rapid climate change and ocean acidification. Science 318, 1737–1742. doi: 10.1126/science.1152509

Khanaev, I. V., Kravtsova, L. S., Maikova, O. O., Bukshuk, N. A., Sakirko, M. V., Kulakova, N. V., et al. (2018). Current state of the sponge fauna (Porifera: Lubomirskiidae) of Lake Baikal: Sponge disease and the problem of conservation of diversity. J. Great Lakes Res. 44, 77–85. doi: 10.1016/j.jglr.2017.10.004

Kolmogorov, M., Armstrong, J., Raney, B. J., Streeter, I., Dunn, M., Yang, F. et al. (2018). Chromosome assembly of large and complex genomes using multiple references. Genome Research 28, 1720–1732.

Kravtsova, L. S., Izhboldina, L. A., Khanaev, I. V., Pomazkina, G. V., Rodionova, E. V., Domysheva, V. M. et al. (2014). Nearshore benthic blooms of filamentous green algae in Lake Baikal. J. Great Lakes Res. 40, 441–448. doi: 10.1016/j.jglr.2014.02.019

Latyshev, N. A., Zhukova, N. V., Efremova, S. M., Imbs, A. B., and Glysina, O. I. (1992). Effect of habitat on participation of symbionts in formation of the fatty acid pool of freshwater sponges of Lake Baikal. Comp. Biochem. Physiol. 102 B, 961–965.

Luter, H. M., Whalan, S., and Webster, N. S. (2010). Exploring the role of microorganisms in the disease-like syndrome affecting the sponge *Ianthella basta*. Appl. Environ. Microbiol. 76, 5736–5744. doi: 10.1128/AEM.00653-10

Matz, C., Webb, J. S., Schupp, P. J., Phang, S. Y., Penesyan, A., Egan, S. et al. (2008). Marine biofilm bacteria evade eukaryotic predation by targeted chemical defense. PLoS ONE 3, e2744. doi: 10.1371/journal.pone.0002744

Miller, A W., and Richardson, L. L. (2011). A meta-analysis of 16S rRNA gene clone libraries from the polymicrobial black band disease of corals. FEMS Microbiol. Ecol. 75, 231–241. doi: 10.1111/j.1574-6941.2010.00991.x

Miller, S. I., and Salama, N. R. (2018). The gram-negative bacterial periplasm: Size matters. PLOS Biology 16, e2004935. doi: 10.1371/journal.pbio.2004935

Nurk, S., Bankevich, A., Antipov, D., Gurevich, A. A., Korobeynikov, A., Lapidus, A., et al. (2013). Assembling single-cell genomes and mini-metagenomes from chimeric MDA products J. Comput. Biol. 20, 714–737. doi: 10.1089/cmb.2013.0084

Olson, J. B., Gochfeld, D. J., and Slattery, M. (2006). *Aplysina red* band syndrome: a new threat to Caribbean sponges. Diseases of Aquatic Organisms 71, 163–168. doi: 10.3354/dao071163

Osinga, R., Armstrong, E., Burgess, J. G., Hoffmann, F., Reitner, J., and Schumann-Kindel, G. (2001). Sponge-microbe associations and their importance for sponge bioprocess engineering. Hydrobiologia 461, 55–62. doi: 10.1023/A:1012717200362

Page, A. J., Cummins, C. A., Hunt, M., Wong, V. K., Reuter, S., Holden, M. T. G., et al. (2015). Roary: Rapid large-scale prokaryote pan genome analysis. Bioinformatics 31, 3691–3693. doi: 10.1093/bioinformatics/btv421

Pantanella, F., Berlutti, F., Passariello, C., Sarli, S., Morea, C., and Schippa, S. (2007). Violacein and biofilm production in *Janthinobacterium lividum*. Appl. Environ. Microbiol. 102, 992–999. doi: 10.1111/j.1365-2672.2006.03155.x

Petrushin, I. S., Belikov, S. I., and Chernogor, L. I. (2019). Draft genome sequence of *Janthinobacterium* sp. strain SLB01, isolated from the diseased sponge *Lubomirskia baicalensis*. Microbiol. Resour. Announc. 8, e01108. doi: 10.3390/ijms21218128

Petrushin, I., Belikov, S., and Chernogor, L. (2020). Cooperative interaction of *Janthinobacterium* sp. SLB01 and *Flavobacterium* sp. SLB02 in the diseased sponge *Lubomirskia baicalensis*. IJMS 21, 8128. doi: 10.3390/ijms21218128

Pita, L., Rix, L., Slaby, B. M., Franke, A., and Hentschel, U. (2018). The sponge holobiont in achanging ocean: from microbes to ecosystems. Microbiome 6, 46. doi: 10.1186/s40168-018-0428-1

Rodrigues, A. L., Gocke, Y., Bolten, C., Brock, N. L., Dickschat, J. S. and Wittmann, C. (2012). Microbial production of the drugs violacein and deoxyviolacein: analytical development and strain comparison. Biotechnol. Lett. 34, 717–720. doi: 10.1007/s10529-011-0827-x.

Silverman, J. M., Brunet, Y. R., Cascales, E., and Mougous, J. D. (2012). Structure and regulation or the type VI secretion system. Annu. Rev. Microbiol. 66, 453–472. doi: 10.1146/annurev-micro-121809-151619

Stabili, L., Cardone, F., Alifano, P., Tredici, S. M., Piraino, S., Corriero, G., et al. (2012). Epidemic mortality of the sponge *Ircinia variabilis* (Schmidt, 1862) associated to proliferation of a *Vibrio bacterium*. Microb. Ecol. 64, 802–813. doi: 10.1007/s00248-012-0068-0

Taylor, M. W., Radax, R., Steger, D., and Wagner, M. (2007) Sponge-associated microorganisms: evolution, ecology, and biotechnological potential. Microbiol. and Mol. Biol. Rev. 71, 295–347. doi: 10.1128/MMBR.00040-06.

Teeseling, M. C. F., Pedro, M. A., and Cava, F. (2017). Determinants of bacterial morphology: from fundamentals to possibilities for antimicrobial targeting. Front. Microbiol. 8. doi.org/10.3389/fmicb.2017.01264

Timoshkin, O. A., Samsonov, D. P., Yamamuro, M., Moore, M. V., Belykh, O. I., Malnik, V.V., et al. (2016). Rapid ecological change in the coastal zone of Lake Baikal (East Siberia): Is the site of the world’s greatest freshwater biodiversity in danger? J. Great Lakes Res. 42, 487–497. doi: 10.1016/j.jglr.2016.02.011

Valdes, N., Soto, P., Cottet, L., Alarcon, P., Gonzalez, A., Castillo, A. et al. (2015). Draft genome sequence of *Janthinobacterium lividum* strain MTR reveals its mechanism of capnophilic behavior. Stand. Genomic Sci. 10, 110. doi: 10.1186/s40793-015-0104-z.

Wang, J., Brodmann, M., and Basler, M. (2019). Assembly and subcellular localization of bacterial type VI secretion systems. Annu. Rev. Microbiol. 73, 621–638. doi: 10.1146/annurev-micro-020518-115420

Waterhouse, R. M., Seppey, M., Simao, F. A., Manni, M., Ioannidis, P., Klioutchnikov, G. et al. (2018). BUSCO applications from quality assessments to gene prediction and phylogenomics. Mol. Biol. and Evol. 35, 543–548. doi: 10.1093/molbev/msx319

Webster, N.S., Negri, A. P., Webb, R. I., and Hill, R. T. (2002). A spongin-boring alpha-proteobacterium is the etiological agent of disease in the Great Barrier Reef sponge *Rhopaloeides odorabile*. Marine Ecology-progress Series 232, 305–309. doi.org/10.3354/meps232305

Webster, N. S., Smith, L. D., Heyward, A. J., Watts, J. E., Webb, R. I., Blackall, L. L. et al. (2004). Metamorphosis of a scleractinian coral in response to microbial biofilms. Appl. Environ. Microbiol. 70, 1213–1221. doi: 10.1128/AEM.70.2.1213-1221.2004

Webster, N. S. (2007). Sponge disease: a global threat? Environ. Microbiol. 9, 1363–1375. doi: 10.1111/j.1462-2920.2007.01303.x.

Webster, N. S., Xavier, J. R.., Freckelton, M., Motti, C. A., and Cobb, R. (2008). Shifts in microbial and chemical patterns within the marine sponge *Aplysina aerophoba* during a disease outbreak. Environ. Microbiol. 10, 3366–3376. doi: 10.1111/j.1462-2920.2008.01734.x.

Webster, N.S., and Taylor, M. W. (2012). Marine sponges and their microbial symbionts: love and other relationships. Environ. Microbiol. 14, 335–346. doi: 10.1111/j.1462-2920.2011.02460.x.

Weisz, J. B., Lindquist, N., and Martens, C.S. (2008). Do associated microbial abundances impact marine demosponge pumping rates and tissue densities? Oecologia 155, 367–376.

